# Chemokine signaling links cell cycle progression and cilia formation for left-right symmetry breaking

**DOI:** 10.1101/578351

**Authors:** Jingwen Liu, Chengke Zhu, Guozhu Ning, Liping Yang, Yu Cao, Sizhou Huang, Qiang Wang

## Abstract

Zebrafish dorsal forerunner cells (DFCs) undergo vigorous proliferation during epiboly and then exit cell cycle to generate Kupffer’s vesicle (KV), a ciliated organ necessary for establishing left-right (L-R) asymmetry. DFC proliferation defects are often accompanied by impaired cilia elongation in KV, but the functional and molecular interaction between cell-cycle progression and cilia formation remains unknown. Here we show that chemokine receptor Cxcr4a is required for L-R laterality by controlling DFC proliferation and KV ciliogenesis. Functional analysis revealed that Cxcr4a accelerates G1/S transition in DFCs and stabilizes Foxj1a, a master regulator of motile cilia, by stimulating Cyclin D1 expression through ERK1/2 signaling. Mechanistically, Cyclin D1-CDK4/6 drives G1/S transition during DFC proliferation and phosphorylates Foxj1a, thereby disrupting its association with Psmd4b, a 19S regulatory subunit. This prevents the ubiquitin-independent proteasomal degradation of Foxj1a. Our study uncovers a role for Cxcr4 signaling in L-R patterning and provides fundamental insights into the molecular linkage between cell-cycle progression and ciliogenesis.

**Author summary:** During the organogenesis of zebrafish L-R organizer named KV, DFCs proliferate rapidly during epiboly and then exit the cell cycle to differentiate into ciliated epithelial KV cells. Cell cycle defects in DFCs are often accompanied by an alteration in KV cilia elongation. However, whether the cell cycle and cilia formation are mechanistically linked remains as an open question. In this study, we report that Cxcr4 signaling is required for DFC proliferation and KV ciliogenesis. We reveal that Cxcl12b/Cxcr4a signaling activates ERK1/2, which then promotes Cyclin D1 expression. Cyclin D1-CDK4/6 accelerates the G1/S transition in DFCs, while also facilitates cilia formation via stabilization of Foxj1a. Notably, Foxj1 undergoes proteasomal degradation via Ub-independent pathway during KV organogenesis. Our study further demonstrates that CDK4 phosphorylates and stabilizes Foxj1a by disrupting its association with Psmd4b, a 19S regulatory subunit. In summary, Cxcl12b/Cxcr4a chemokine signaling links cell cycle progression and cilia formation for L-R symmetry breaking via regulating Cyclin D1 expression.

## Introduction

Vertebrates exhibit striking left-right (L-R) asymmetries in the structure and position of their cardiovascular and gastrointestinal systems. Initially, early embryos develop symmetrically along the prospective body midline. This embryonic symmetry is broken during somite stages when an asymmetric fluid flow is generated by motile cilia within the L-R organizer (LRO), a transient structure located at the posterior end of the notochord [1]. Specifically, in zebrafish, the ciliated LRO is referred to as Kupffer’s vesicle (KV), which forms from dorsal forerunner cells (DFCs), a group of superficial cells in the organizer region of the gastrula [2,3]. It has been well established that the architecture of KV cells and asymmetric KV cilia generate a counter-clockwise nodal flow. This leads to the asymmetrical expression of early laterality genes, including *nodal-related southpaw* (*spaw*) and *pitx2c* in the left lateral plate mesoderm (LPM), and ultimately the establishment of L-R asymmetric patterning [4]. The origin of L-R asymmetry is conserved across many vertebrates, and defects in the establishment of these asymmetries can result in a broad spectrum of birth defects, often including congenital heart malformations [5,6].

The progression of cells through the G1 and S phases of the cell cycle is tightly controlled by the sequential activation of a family of serine-threonine kinases known as the cyclin-dependent kinases (CDKs). CDK4 and its homologous CDK6 are activated by D-type cyclins in early to mid-G1 phase, whereas CDK2 is activated by E- and A-type cyclins during late G1 and S phase, respectively [7]. Recent evidence indicates that cell cycle dynamics have emerged as a key regulator of stem cell fate decisions [8–10]. Specifically, cyclin D proteins have been shown to activate CDK4/6, which restricts the activity of Smad2/3 in late G1 phase and results in a switch from endoderm to neuroectoderm potential in human pluripotent stem cells [11]. The G1 cyclin proteins together with their associated CDKs also play essential, direct roles in the maintenance of cell stemness and in the regulation of cell fate specification in mouse embryonic stem cells by phosphorylation and stabilization of the core pluripotency factors, Nanog, Sox2, and Oct4 [12]. In addition, CDK4/CyclinD1 overexpression has been shown to prevent G1 lengthening and functions to inhibit neurogenesis in mouse embryos [13]. In zebrafish, DFCs vigorously proliferate and collectively migrate towards the vegetal pole during epiboly stages. They then cluster and differentiate into polarized epithelial cells of KV [4,14,15]. Interestingly, DFC proliferation defects are often accompanied by impaired cilia elongation in KV [16], indicating a possible connection between cell-cycle events and KV cilia formation. However, the underlying mechanism remains poorly understood.

Foxj1, a forkhead domain-containing transcription factor that is expressed in various ciliated tissues, has been associated with motile cilia formation and L-R axis development in mammals [17,18]. In zebrafish, two *foxj1* paralogs have been identified, including *foxj1a* and *foxj1b* [19]. *foxj1a* has been shown to be highly expressed in the DFCs toward the end of gastrulation and plays a primary role in KV ciliogenesis, while *foxj1b* is expressed in the otic vesicle where it has been shown to regulate motile cilia formation [19,20]. The expression level of *foxj1a* transcripts has been shown to be regulated by the Hedgehog, Wnt/β-catenin and FGF signalling pathways [19,21,22]. Lnx2b, a RING domain containing E3 ubiquitin (Ub) ligase, which is specifically expressed in the migratory DFCs and developing KV, plays a critical role in the establishment of L-R laterality. This indicates the involvement of protein ubiquitination in the determination of L-R asymmetry [23]. However, whether the function of Foxj1 protein in KV ciliogenesis is regulated by Ub modification remains unknown.

Chemokines are small (8-14 kDa) vertebrate-specific proteins that can be categorized into four subgroups according to the presence and position of conserved cysteine residues (C, CC, CXC, and CX3C) [24]. Among chemokines of the CXC class, the stromal cell-derived factor 1 (SDF-1/CXCL12) and its receptor CXCR4, which were first identified due to their primary role in leukocyte homing, have been implicated in the regulation of cell adhesion and migration during embryonic development [24–26]. Interestingly, in zebrafish, two Cxcl12 ligands and two Cxcr4 receptors were found to be expressed across a wide range of cell types and developmental stages, and were found to act as discrete pairs to direct cell migration [25]. Cxcl12a-Cxcr4b signaling controls processes such as the directional migration of primordial germ cells, the collective migration of the lateral line primordium, and the formation of the truck lymphatic network [27–29]. On the other hand, the Cxcl12b-Cxcr4a axis has been shown to play a role in endodermal morphogenesis, vascular system patterning, and the migration and prechondrogenic condensation of cranial neural crest cells [30–33]. It has been shown previously that *cxcr4a* and *cxcr4b* possess mutually exclusive expression patterns in the majority of cell lineages [34]. For example, *cxcr4a* but not *cxcr4b* is expressed in the primordium of KV [34]. While *cxcr4b* expression reveals an asymmetric pattern in habenular neurogenesis, the *cxcr4b* mutant *odysseus* displays no obvious phenotype in L-R epithalamic asymmetry [35]. These observations bring into question whether the signaling cascades initiated by Cxcl12b and Cxcr4a play a role in the establishment of L-R asymmetry.

Here, we provide evidence suggesting that the Cxcl12b-Cxcr4a axis is essential for L-R asymmetric development. *Cxcr4a^um20^* mutants were found to exhibit poor DFC proliferation and abnormal KV cilia formation. Specifically, depletion of *cxcr4a* in DFCs was found to lead to a significant decrease in ERK1/2 signal activation, which was essential for the expression of *cyclin D1*. Subsequent biochemical and functional approaches demonstrated that Cyclin D1-CDK4/6 functions to accelerate the G1/S transition to promote DFC proliferation and stabilize Foxj1a for cilia formation. Mechanistically, CDK4 phosphorylates zFoxj1a at T102 and then disrupts its association with Psmd4b, which in turn prevents the ubiquitin-independent proteasomal degradation of Foxj1a protein. Therefore, Cxcl12b/Cxcr4a chemokine signaling links cell cycle progression and cilia formation for L-R symmetry breaking via regulating Cyclin D1 expression.

## Results

### Cxcl12b-Cxcr4a axis is required for L-R laterality

*cxcr4a* was previously found to be expressed in KV progenitors at the end of gastrulation [34]. To address the detailed expression patterns of *cxcr4a*, whole mount *in situ* hybridization was carried out during early zebrafish embryogenesis. As shown in S1A Fig, *cxcr4a* expression was observed in endoderm cells and migrating DFCs throughout gastrulation. At early somite stages, concomitant with the onset of anterior neural plate expression, *cxcr4a* was also found to be activated in the developing KV cells besides the central lumen (S1B Fig). Therefore, we hypothesized that *cxcr4a* might play a critical role in KV organogenesis and L-R asymmetric patterning.

We then set out to test whether *cxcr4a* is required for L-R development in *cxcr4a^um20^* embryos carrying loss-of-function mutations in *cxcr4a* gene, which lead to defects in lateral dorsal aorta formation (S1C Fig) [33]. In order to analyze the laterality information in homozygous *cxcr4a^um20^* mutants, we examined cardiac development by WISH against cardiac myosin light chain 2 (*cmlc2*). At 48 hours post fertilization (hpf), the majority of wild-type embryos showed a heart tube looping to the right (D-loop) (Fig 1A and 1B). However, the heart localization became randomized in *cxcr4a^um20^* mutants, of which 21% showed a “no-looping” or reversed “left-looping” heart (L-loop) (Fig 1A and 1B). In addition, in *cxcr4a^um20^* embryos, the livers were also observed to be randomized in laterality as revealed by *hhex* expression (Fig 1C and 1D).

**Fig 1.**
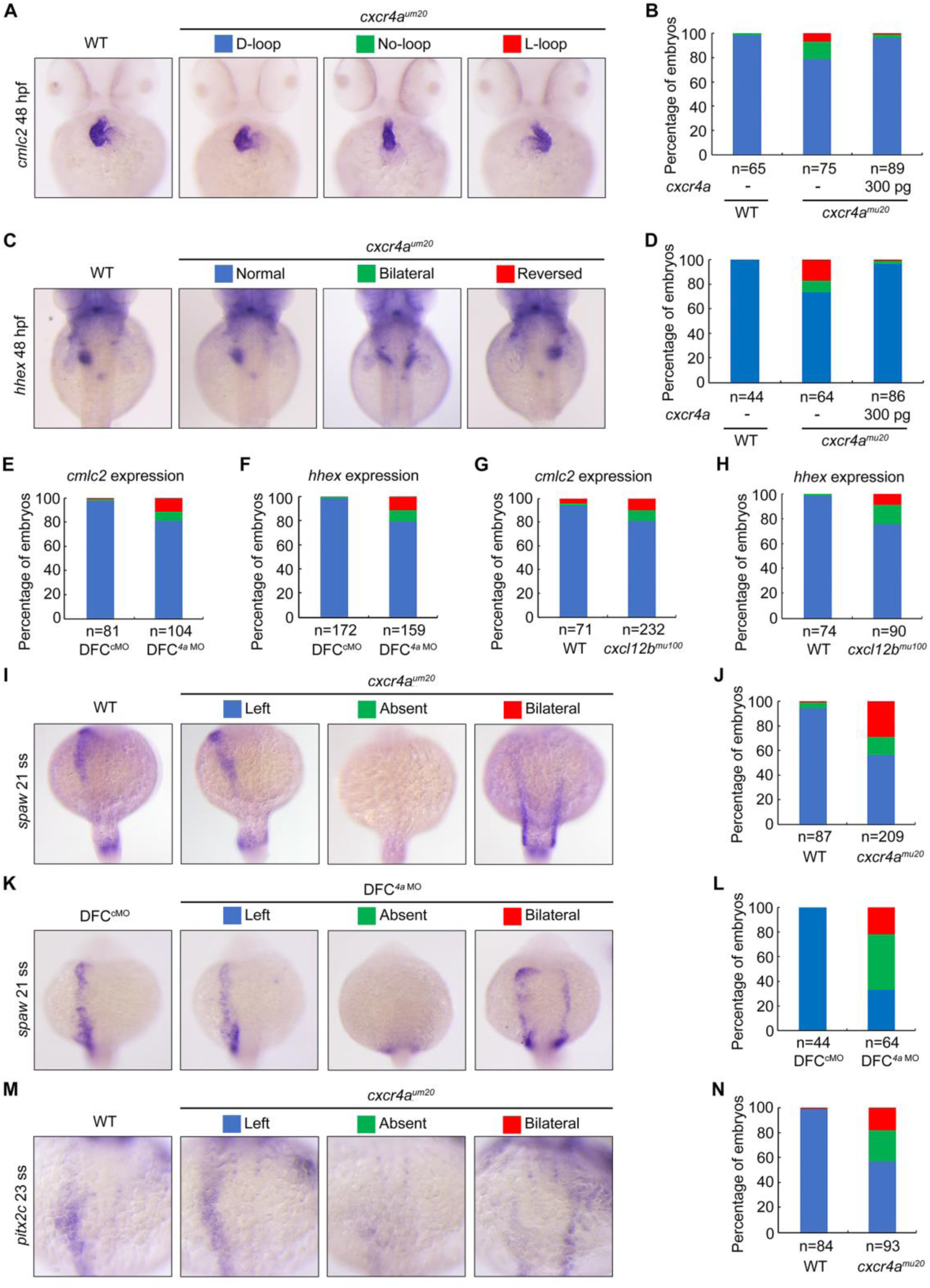
Cxcl12b-Cxcr4a signaling axis is essential for L-R asymmetric development. (A-D) Wild-type embryos and *cxcr4a^mu20^* mutants injected with or without 300 pg *cxcr4a* mRNA at the 256-cell stage were examined for cardiac looping and liver laterality at 48 hpf by WISH against *cmlc2* (a) and *hhex* (C). Embryos with different phenotypes were shown in ventral (A) or dorsal view (C). Statistical data were shown in (B) and (D). (E-H) Statistical data for the expression patterns of *cmlc2* and *hhex* at 48hpf in wild-type embryos injected with 8 ng *cxcr4a* MO (4a MO) at the 256-cell stage (E and F) and *cxcl12b* mutants (G and H). (I-N) *cxcr4a* deficiency alters Nodal gene expression pattern. Representative images of *spaw* and *pitx2c* expression in *cxcr4a* mutants (I and M) and morphants (K). All embryos were shown in dorsal views with anterior on the top. Ratios of embryos were shown in (J, L and N).

Because *cxcr4a* depletion would impair endoderm cell migration during gastrulation and cause bilateral duplication of endodermal organs such as the liver [30], we injected a previously validated morpholino that targets *cxcr4a* (*4a* MO) into embryonic yolk at the mid-blastula stage (256-cell stage) to specifically block *cxcr4a* activity in DFC/KV cells, as demonstrated previously [36]. In comparison to injection with a standard control morpholino (cMO), injection of *4a* MO led to similar laterality abnormalities as observed in *cxcr4a^um20^* mutants (Fig 1E and 1F). This suggests that the organ localization defects are not secondary effects of impaired endoderm migration. Therefore, the *cxcr4a* expression in DFC/KV cells is required for L-R laterality. In addition, the deficiency of *cxcl12b* in *cxcl12b^mu100^* mutants [37], was also found to result in laterality defects (Fig 1G and 1H), indicating that the Cxcl12b-Cxcr4a signaling pathway is critical for L-R symmetry breaking.

Because organ laterality is regulated by evolutionally conserved asymmetric L-R gene expression in vertebrates, we next examined the expression patterns of *spaw* and its downstream gene *pitx2c* [1,38]. At the late somite stages, we observed *spaw* and *pitx2c* expression in the left LPM in wild-type embryos, whereas expression of these genes was found to be bilateral or absent in *cxcr4a^um20^* mutants and DFC*^4a^* ^MO^ embryos (Fig 1I-1N). Interestingly, the bilateral expression domain of *spaw* in a subset of *cxcr4a^um20^* mutants was located in the more posterior region in the LPM (Fig 1I), indicating a delay in the anterior spreading of *spaw* expression.

Collectively, these results demonstrate a sustaining expression of *cxcr4a* in the DFC/KV cells and implicate a crucial role of Cxcl12b-Cxcr4a chemokine signaling in L-R laterality determination.

### Ablation of *cxcr4a* compromises KV organogenesis and ciliogenesis

To determine whether a loss of *cxcr4a* alters KV morphogenesis, we first examined the formation of DFC clusters during gastrulation in *cxcr4a^um20^* mutants carrying the transgenic DFC/KV reporter *sox17:GFP*. We observed that, in both wild-type embryos and *cxcr4a^um20^* mutants, the GFP-positive DFCs were maintained as a cohesive group and migrated towards the vegetal pole during mid- to late-gastrulation (S2A Fig). Meanwhile, in comparison to control embryos, *cxcr4a^um20^* mutants exhibited a normal expression pattern of *sox17* transcripts in the DFC clusters (S2B Fig). Based on these observations, we concluded that *cxcr4a* is unnecessary for the specification, clustering, and collective migration of DFCs.

We then examined the morphology of KV in live embryos at the 10-somite stage, at which point KV was well formed [4]. Wild-type embryos exhibited a normal button-like KV at the terminus of the notochord, as observed under bright-field microscopy (Fig 2A). In contrast, a large majority of *cxcr4a^um20^* mutants displayed a smaller or tiny/absent KV (Fig 2A and 2B). In order to monitor the dynamic changes during KV formation, we carried out *in vivo* time-lapse image analysis on *cxcr4a*-deficient *Tg(sox17:GFP)* embryos from the 1- to 6-somite stages. DFCs were found to have already rearranged into a single rosette concurrent with the formation of the preliminary lumen in 1-somite stage wild-type and *cxcr4a^um20^* mutant embryos (Fig 2C). However, the GFP-positive KV appeared to be dramatically smaller in *cxcr4a^um20^* embryos in comparison to control animals from the 3- to 6-somite stages (Fig 2C). We next looked into the apical-basal polarity of KV epithelial cells, which is critical for the correct establishment of L-R asymmetry [39]. Immunostaining experiments revealed that the distributions of the basal-lateral marker E-cadherin and the apical marker aPKC in KV epithelial cells of 10-somite stage *cxcr4a^um20^* embryos were correct (S2C-S2D Fig). These observations suggest that *cxcr4a* is critical for organ size control, but is not required for epithelial cell polarization during KV organogenesis.

**Fig 2.**
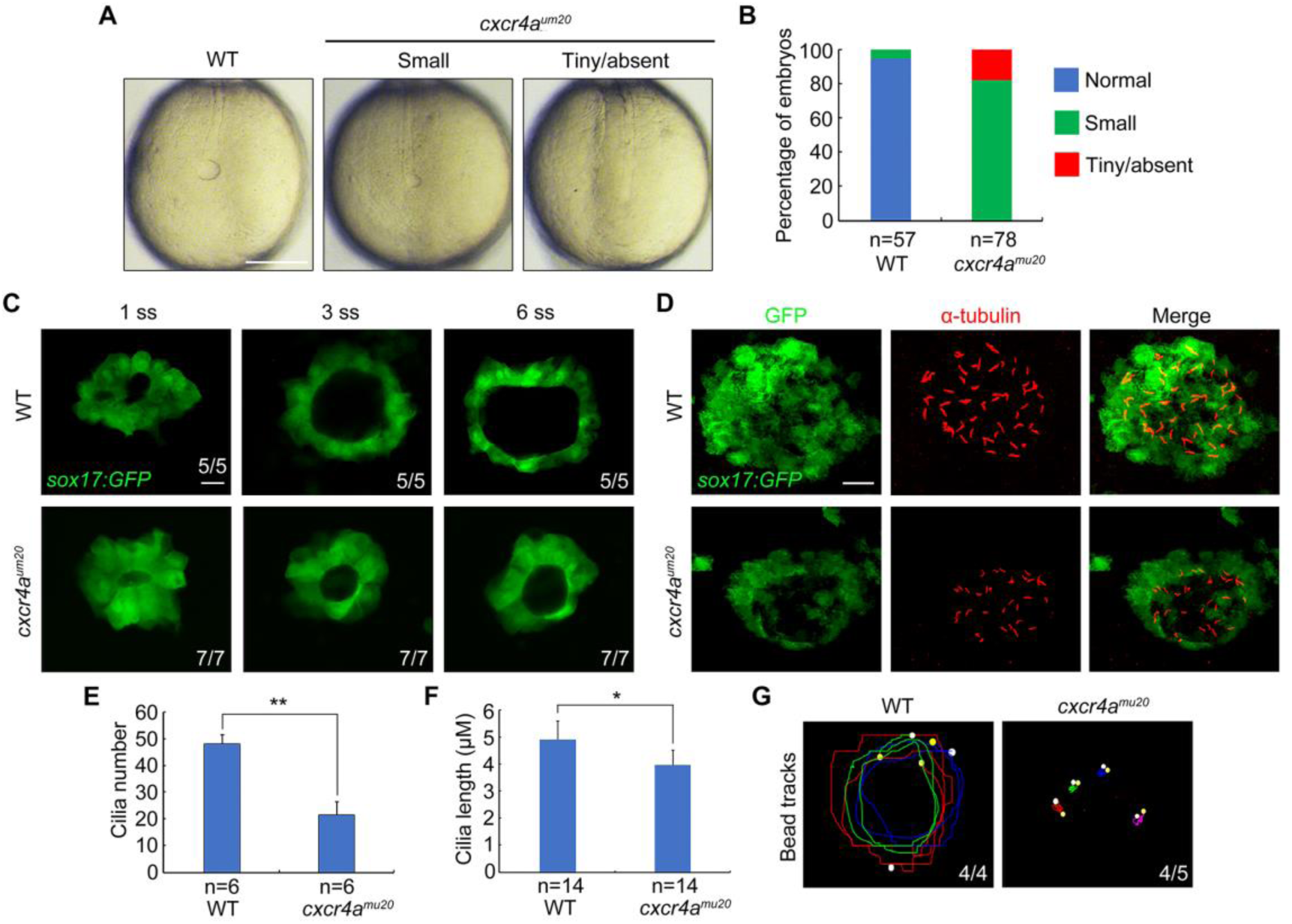
*cxcr4a* is indispensable for KV formation and ciliogenesis. (A-B) Light micrographs at the 10-somite stage showed smaller or even absent KVs in *cxcr4a^mu20^* mutants. Scale bar, 200 μm. Embryo ratios with different KV sizes were shown in (B). (C) Time-lapse confocal images from the 1-somite stage to the 6-somite stage showed the dynamic formation of KV in wild-type and *cxcr4a* deficient *Tg(sox17:GFP)* embryos. Scale bar, 20 μm. The ratios of affected embryos was indicated. (D-F) Fluorescent immunostaining of KV using anti-GFP and anti-acetylated Tubulin antibodies at the 10-somite stage in wild-type embryos and *cxcr4a^mu20^* mutants. Scale bar, 20 μm. Cilia average number and length were quantified from three independent experiments and the group values were expressed as the mean ± SD (E and F). Student’s *t*-test, \**P*<0.05, \*\**p* < 0.01. (G) Fluorescent bead tracking experiments showed that fluorescent beads moved in a persistent counter-clockwise fashion in wild-type embryos, but had no directional flow in *cxcr4a^mu20^* mutants. White spots, yellow spots, and curves mark the start points, the end points, and the tracks of bead movements, respectively.

Monocilia in KV are known to generate a counter-clockwise fluid flow, which creates asymmetrical signals required to break L-R symmetry [4]. We next analyzed KV cilia formation by probing for acetylated tubulin (α-Tubulin) in the *cxcr4a^um20^* embryos at the 10-somite stage. We found that, in comparison with control embryos, *cxcr4a^um20^* mutants exhibited a significant decrease in cilia number and a steady reduction in cilia length (Fig 2D-2F). We then sought to determine whether the KV directional fluid flow was altered in *cxcr4a^um20^* mutants. Fluorescent beads were injected into KVs at the 6-somite stage and the movements of the beads were tracked at the 10-somite stage. The fluorescent beads moved in a persistent counter-clockwise fashion in wild-type embryos, whereas they exhibited no directional flow in *cxcr4a*-deficient embryos (Fig 2G; S1-S2 Videos). Therefore, these results indicated that *cxcr4a* is indispensable for KV ciliogenesis and cilia-driven fluid flow.

### Absence of *cxcr4a* attenuates G1/S transition and zFoxj1a protein expression

Our studies suggest that *cxcr4a* deficiency leads to smaller KV size as well as fewer KV cilia. Interestingly, the majority of the KV epithelial cells in *cxcr4a* mutants exhibited an intact apical-basal polarity and formed notably shortened cilia (Fig 2D and S3C-S3D Fig). This suggests that the altered KV cilia numbers may be caused by defects in cell proliferation. To address this issue, we first examined the proliferation profile of DFC/KV cells by performing bromodeoxyuridine (BrdU) incorporation assays in *Tg(sox17:GFP)* embryos during gastrulation and early somite stages. Consistent with previous reports [16], approximately 70% of DFCs were positively stained with BrdU at the mid-gastrulation stage, whereas very few BrdU-positive cells were observed in the developing KV at the bud and the 6-somite stages (Fig 3A and 3B), suggesting that vigorous proliferation occurs in DFCs during epiboly stages and then declines at the end of gastrulation. Impressively, we found a dramatic decrease of the BrdU-positive DFC number in *cxcr4a*-deficient embryos at mid-gastrulation stage (Fig 3C and 3D), indicating a crucial requirement of *cxcr4a* in DFC proliferation.

**Fig 3.**
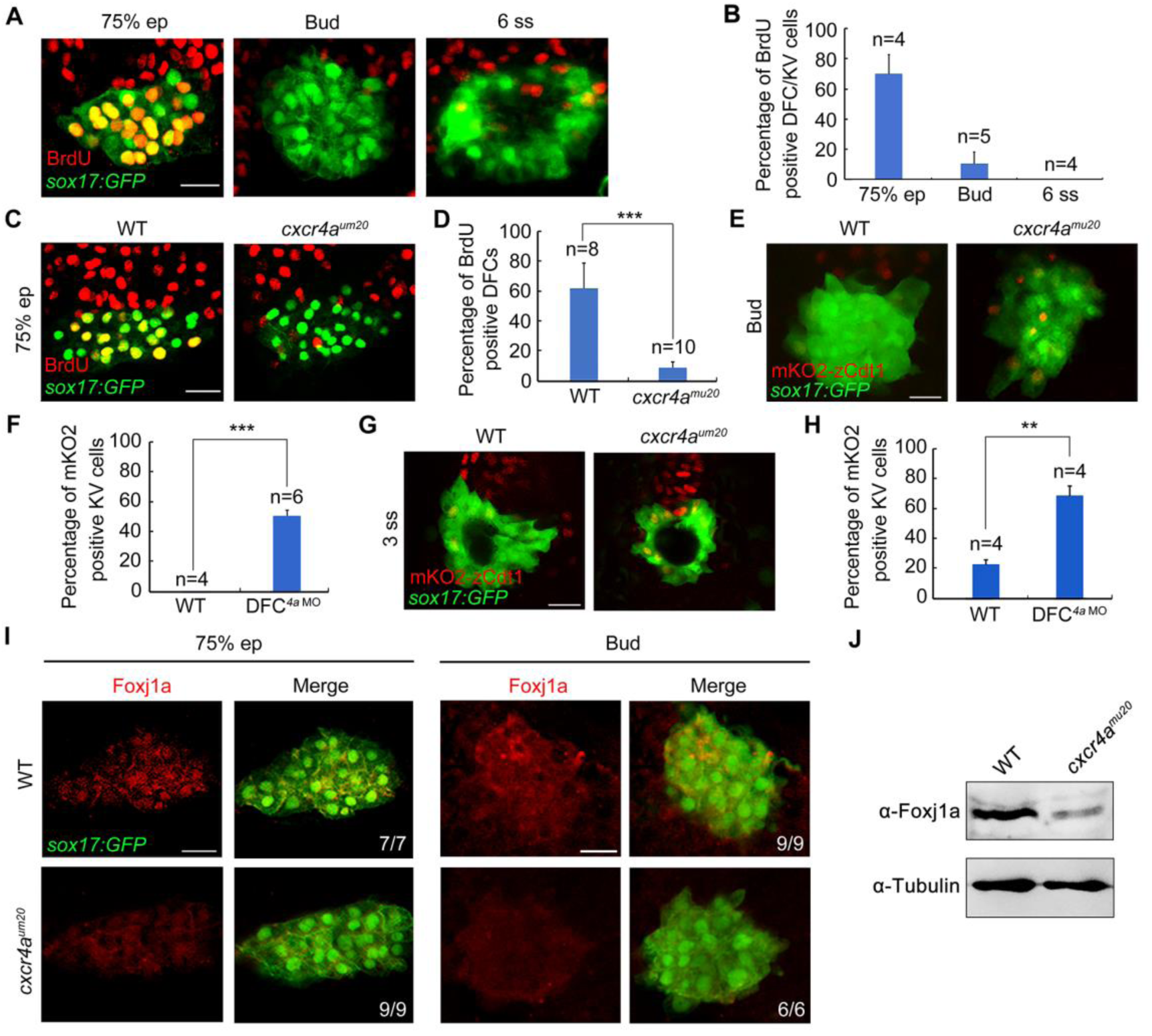
Depletion of *cxcr4a* impairs G1/S transition and Foxj1a protein expression in DFCs. (A-B) Representative confocal sections of BrdU-positive DFCs and KV cells at the indicated stages (A). Dorsal views with anterior on the top. Scale bar, 20 μm. The percentage of BrdU-positive cells among GFP-positive DFCs and KV cells were quantified from the indicated embryo numbers in three independent experiments and the group values are expressed as the mean ± SD (B). (C-D) BrdU incorporation experiments showed reduced proliferating DFCs in *cxcr4a^mu20^* mutants at the 75% epiboly stage. Scale bar, 20 μm. Statistical data from three independent experiments was shown in (D). Student’s *t-*test, \*\*\**P*<0.001. (E-H) Depletion of *cxcr4a* inhibits the G1/S transition in DFCs. Representative confocal sections of wild-type and *cxcr4a*-deficient *Tg(sox17:GFP;EF1α:mKO2-zCdt1(1/190))* embryos at the bud and 3-somite stages were shown in (E and G). Dorsal views with anterior to the top. Scale bar, 20 μm. The percentage of mKO2-positive KV cells were quantified from three independent experiments (F and H). The significance of differences compared with the wild-type group were analyzed with the Student’s *t*-test, \*\**p* < 0.01; \*\*\**p* < 0.001. (I-J) *cxcr4a* deficiency downregulates zFoxj1a protein expression levels. Wild-type and *cxcr4a*-deficient *Tg(sox17:GFP)* embryos were harvested at the 75% epiboly and bud stages, and then subjected to immunostaining (I) and western blot analysis (J) with the indicated antibodies. Scale bar, 20 μm.

We next examined the detailed effects of *cxcr4a* deficiency on DFC cycle progression in *Tg(sox17:GFP;EF1α:mKO2-zCdt1(1/190))* double transgenic embryos, a model in which cells in G1 phase exhibit red nuclear fluorescence [40]. Because the G1/S transition is very short in cells which undergo rapid mitotic cycles [40], we were unable to identify any mKO2-zCdt1-positive DFCs in both control embryos and *cxcr4a^um20^* mutants during gastrulation. Interestingly, while the majority of cells in the developing KV enter into a quiescent state at the end of gastrulation (Fig 3A and 3B), we observed no or only few KV progenitors with mKO2-zCdt1 fluorescence (Fig 3E-3H). These results, combined with the previous observation that the mKO2-zCdt1 signal was highlighted in differentiated cells, including postmitotic neurons and muscle cells after the 10-somite stage [40], imply that there exist limitations in the ability to dissect cell-cycle behavior using this fluorescent indicator in early zebrafish embryos. However, *cxcr4a* depletion resulted in a marked proportion of KV progenitors with robust mKO2-zCdt1 expression, indicating impaired G1/S transition and an apparent lengthening of the G1 phase (Fig 3E-3H). Nevertheless, our results suggest that Cxcr4a-mediated chemokine signaling is responsible for driving DFC proliferation by accelerating the G1/S transition. This provides an explanation for the smaller KV size which was observed in *cxcr4a^um20^* mutants.

Because zebrafish Foxj1a (zFoxj1a) is a master regulator of KV ciliogenesis [19,20], we then sought to determine whether zFoxj1a expression was affected in *cxcr4a^um20^* mutants. *In situ* hybridization analysis demonstrated normal expression levels of *zfoxj1a* transcripts in *cxcr4a^um20^* embryos during gastrulation (S3 Fig). In contrast, expression levels of zFoxj1a proteins were clearly decreased in *cxcr4a^um20^* mutants as revealed by immunostaining and western blot experiments (Fig 3I and 3J). These analyses provide strong evidence that Cxcr4a signaling is responsible for controlling KV ciliogenesis through regulation of zFoxj1a protein expression at the post-transcription level.

### Cxcr4a-ERK1/2 cascade controls DFC proliferation by regulating cyclin D1 expression

The Cxcl12-Cxcr4 axis is known to regulate cell-cycle progression through GSK-3β/β-catenin and ERK1/2 signaling pathways [41–43]. To determine which of these candidate pathways mediates Cxcr4-regulated DFC proliferation, *cxcr4a^um20^* mutant embryos were immunostained with antibodies against β-catenin or phosphorylated ERK1/2 (p-ERK1/2) at the 75%-epiboly stage. No obvious changes in the cellular distribution of endogenous β-catenin in DFCs were observed in *cxcr4a^um20^* mutants (S4 Fig), indicating that GSK-3β/β-catenin signaling is not altered with *cxcr4a* depletion. However, we found a robust expression of p-ERK1/2 in wild-type DFCs, which was nearly abolished in *cxcr4a-*deficient cells (Fig 4A). Strikingly, DFC-specific overexpression of MEK1^S219D, S223D^, a constitutively activated version of MEK1 (caMEK1) [44], rescued the L-R defects in *cxcr4a^um20^* mutants in a dose-dependent manner (Fig 4B-4E). Therefore, these results demonstrate a role for ERK1/2 signaling downstream of Cxcr4 in organ laterality.

**Fig 4.**
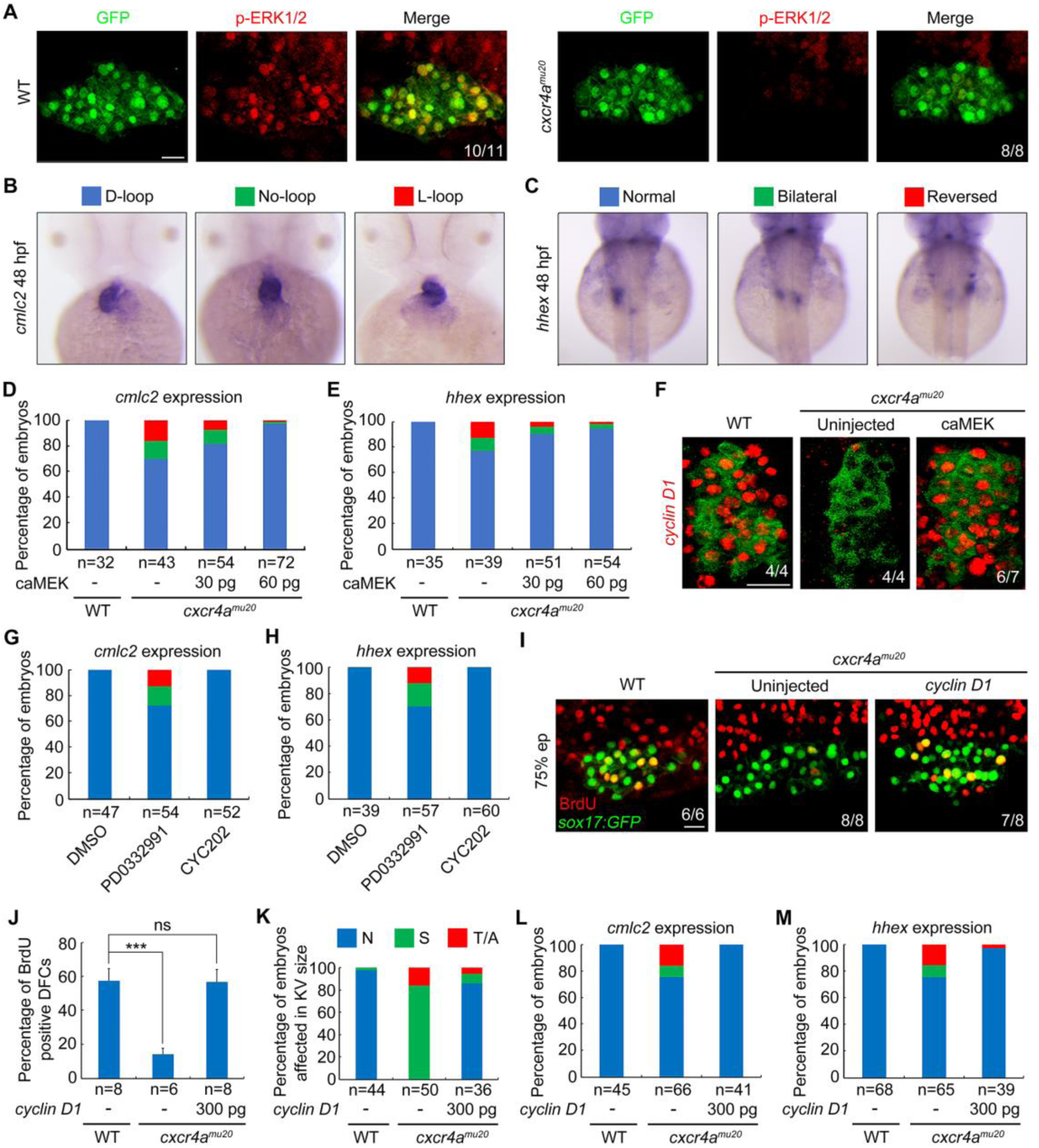
Cxcr4 promotes Cyclin D1 expression through ERK signaling during DFC proliferation. (A) ERK1/2 phosphorylation levels were dramatically decreased in *cxcr4a^mu20^* mutants. Wild-type and *cxcr4a*-deficient *Tg(sox17:GFP)* embryos were harvested at the 75% epiboly stage and subjected to immunostaining for p-ERK1/2 (red) and GFP (green). All embryos were shown in dorsal views with anterior to the top. Scale bar, 20 μm. (B-E) caMEK mRNA overexpression in DFCs rescued L-R patterning defects in *cxcr4a^mu20^* mutants. Different types of heart looping and liver laterality at 48 hpf in *cxcr4a^mu20^* mutants following mid-blastula injection of different caMEK mRNA doses were visualized by *cmlc2* and *hhex* expression (B and C). Quantitative data were shown in (D and E). (F) *Cxcr4a*-deficient *Tg(sox17:GFP)* embryos were injected with 60 pg caMEK mRNA at the 256-cell stage, and then harvested at the 75% epiboly stage for fluorescence *in situ* hybridization experiments with *cyclin D1* (red) and GFP (green) probes. Dorsal views with anterior to the left. Scale bar, 20 μm. (G-H) Wild-type embryos were treated 0.5 μM PD0332991 or 0.2 μM CY202 from the shield stage to bud stage, and then analyzed for L-R patterning defects at 48 hpf by *in situ* hybridizations with *cmlc2* and *hhex* probes. The proportion of treated embryos exhibiting each type of heart looping and liver laterality were shown in (G) and (H). (I-J) Reintroduction of *cyclin D1* into DFCs relieves DFC proliferation defects in *cxcr4a^um20^* mutants. *Cxcr4a*-deficient *Tg(sox17:GFP)* embryos were injected with or without 300 pg *cyclin D1* mRNA at the 256-cell stage, followed by coimmunostaining with anti-BrdU (red) and anti-GFP (green) antibodies at the 75% epiboly stage. Representative images were shown in (I) and the percentage of BrdU-positive DFCs were indicated in (J). Scale bar, 20 μm. Student’s *t-*test, \*\*\**P*<0.001. ns, no significant difference.(K-M) Statistical data shows that DFC-specific overexpression of *cyclin D1* rescued the defects of KV formation (K) and L-R patterning (L and M) in *cxcr4a* mutants.

Among the cell-cycle-regulatory genes, Cyclin D1 expression is known to be specifically activated by the Cxcr4a-ERK1/2 cascade to promote cell proliferation [41,42,45]. Consistent with these previous studies, double fluorescence *in situ* hybridization analyses indicated a dramatic reduction in *cyclin D1* expression in *cxcr4a-*deficient DFCs (Fig 4F). In addition, the introduction of caMEK mRNA counteracted the *cxcr4a* depletion effects on *cyclin D1* expression (Fig 4F). Cell cycle progression from G1 to the S phase is governed by CDK4/6 and CDK2, which are activated by D-type and E- or A-type cyclins, respectively [7]. Interestingly, upon exposure of wild-type embryos to PD0332991, a selective CDK4/6 inhibitor [46], from the shield stage to the bud stage, the resulting animals exhibited L-R defects similar to those observed in *cxcr4a^um20^* mutants (Fig 4G and 4H). At the same time, embryos treated with CYC202, a selective CDK2 inhibitor [47], showed no significant changes in laterality development (Fig 4G and 4H). These results suggest that Cyclin D1-CDK4/6 complexes play critical roles in DFC proliferation and L-R asymmetric development. Importantly, reintroduction of *cyclin D1* into DFCs was found to relieve the inhibition of cell proliferation, the reduction of KV size, and the defects of organ laterality in *cxcr4a^um20^* mutants (Fig 4I-4M). Collectively, our data indicate that the Cxcr4a-ERK1/2 cascade functions in DFC proliferation through regulation of Cyclin D1 expression during zebrafish L-R development.

### CDK4 and its kinase activity is required for Foxj1 protein stabilization

Because the injection of *cyclin D1* mRNA into DFCs was found to rescue the L-R defects in *cxcr4a^um20^* mutants (Fig 4L and 4M), we hypothesized that Cyclin D1 acts downstream of Cxcr4a signaling to control both cell proliferation and cilia formation. In support of this hypothesis, upon injection of 300 pg *cyclin D1* mRNA into mid-blastula stage *cxcr4a^um20^* embryos, we observed that both the number and the length of KV cilia were restored (Fig 5A-5C), indicating a role of Cyclin D1 in ciliogenesis. It is surprising that the DFC-specific delivery of *cyclin D1* mRNA was also able to restore the expression of endogenous Foxj1a protein in *cxcr4a^um20^* embryos (Fig 5D).

**Fig 5.**
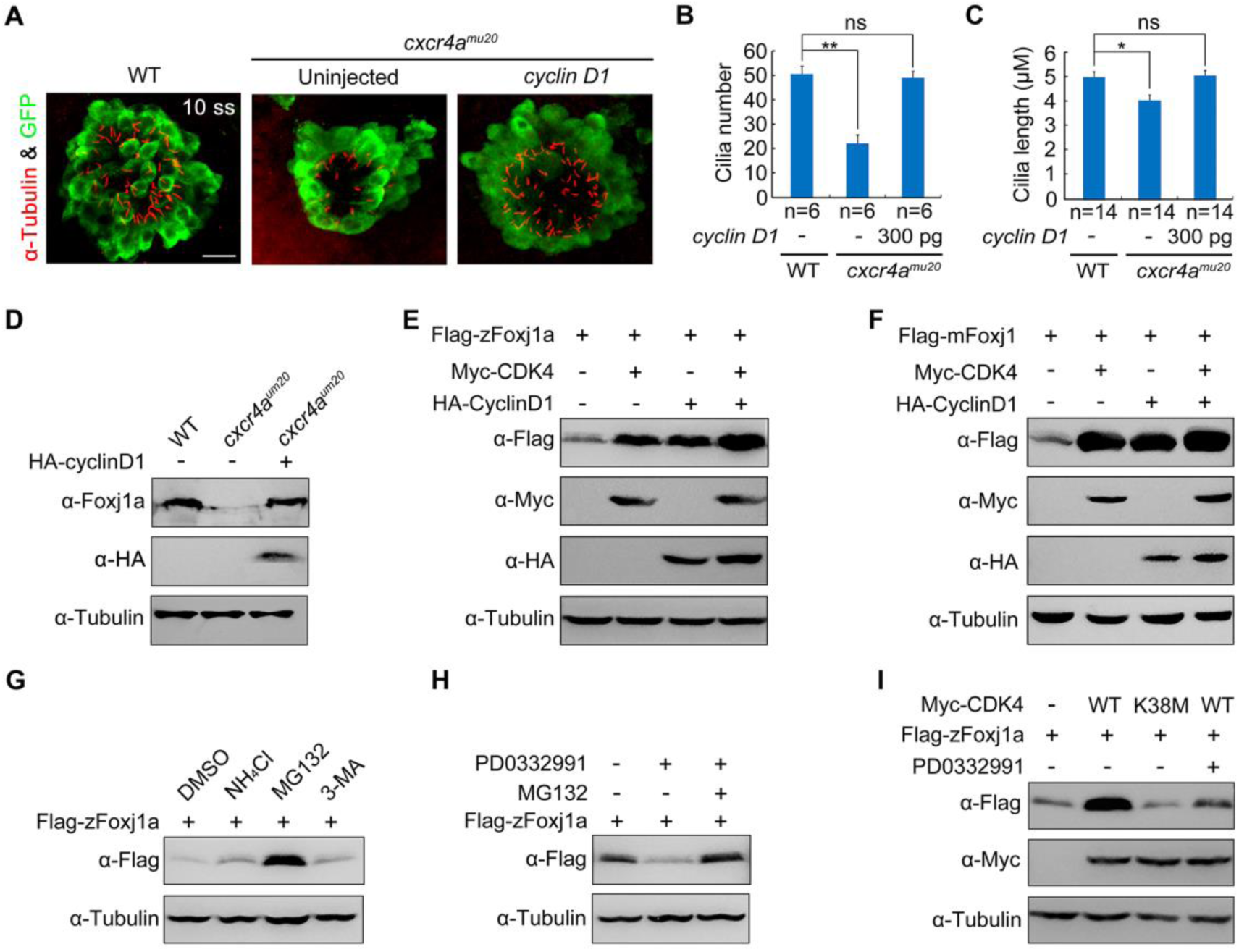
CDK4 stabilizes Foxj1 through its kinase activity. (A-C) Confocal images of 10-somite-stage *cxcr4a*-deficient *Tg(sox17:GFP)* embryos injected with or without 300 pg *cyclin D1* mRNA at the 256-cell stage (A). KV cells were labelled with antibodies against GFP (green) and cilia were visualized by acetylated tubulin immunofluorescence (red). Scale bar, 20 μm. Cilia number (B) and length (C) were analyzed. Student’s *t-*test, \**P*<0.05, \*\**P*<0.01. ns, no significant difference. Note that when *cyclin D1* mRNA was injected into *cxcr4a*-deficient DFCs, both the KV cilia number and length were restored. (D) Western blots of total lysates from 75% epiboly-stage wild-type embryos and *cxcr4a^um20^* mutants injected with or without 300 pg *HA-cyclin D1* mRNA at the 256-cell stage. Tubulin was used as the loading control. Endogenous zFoxj1a protein levels in *cxcr4a^um20^* mutants was rescued by DFC-specific overexpression of HA-cyclin D1. (E-F) CDK4 overexpression alone or together with Cyclin D1 results in an obvious increase in zFoxj1a (E) or mFoxj1 (F) expression. HEK293T cells were transfected with the indicated plasmids. Lysates were analyzed by western blot using the indicated antibodies. (G) HEK293T cells transfected with plasmids encoding Flag-zFoxj1a were treated with the lysozyme inhibitor NH4Cl (20 mM) or the proteasomal inhibitor MG132 (20 μM) or the autophagy inhibitor 3-MA (5 mM) for 5 hours prior to harvest for immunoblotting. (H) Lysates from Flag-zFoxj1a-expressing HEK293T cells treated with CDK4/6 inhibitor PD0332991 (0.5 μM) alone or in combination with MG132 (20 μM) were subjected to immunoblotting. (I) Overexpression of wild-type CDK4 but not its kinase mutant stabilizes zFoxj1a protein. HEK293T cells were transfected with the indicated plasmids. PD0332991 (0.5 μM) was added 5 hours before harvest. Note that PD0332991 treatment blocked CDK4-induced zFOxj1a stabilization.

We next aimed to understand whether Foxj1 protein stability is regulated by Cyclin D1-CDK4/6 complexes. As depicted in Fig 5E, Cyclin D1 or CDK4 overexpression in HEK293T cells notably increased zFoxj1a expression. This increase in exogenously expressed zFoxj1a was even more apparent when Cyclin D1-CDK4 complexes were ectopically expressed (Fig 5E). We noted that Cyclin D1-CDK4 complexes also showed similar effects on the expression levels of mouse Foxj1 (mFoxj1) (Fig 5F). In combination with our observation of unchanged *foxj1a* transcript expression in *cxcr4a^um20^* mutants, these results suggest that Cyclin D1-CDK4 complexes may play crucial roles in the prevention of Foxj1a protein degradation. Indeed, we observed that the treatment of MG132, a proteasome inhibitor, but not NH4Cl, a lysosome inhibitor, or 3-methyladenine (3-MA), a well-characterized inhibitor of autophagy, dramatically stabilized zFoxj1a protein (Fig 5G). In contrast, blocking endogenous CDK activity with PD0332991 treatment was found to result in a clear reduction of zFoxj1a expression (Fig 5H). In addition, the PD0332991 treatment-induced zFoxj1a turnover was completely suppressed by co-treatment with MG132 (Fig 5H). Therefore, Cyclin D1-CDK4 complexes contribute to Foxj1 protein stabilization. To further determine whether CDK4 kinase activity is important for Foxj1 protein stability, we generated a zebrafish kinase deficient mutant denoted CDK4-K38M, in which the ATP-binding site (Lys-38) in the catalytic subunit was mutated to a methionine residue [48]. As shown in Fig 5I, ectopic expression of CDK4-K38M had no effect on the zFoxj1a expression and PD0332991 treatment eliminated CDK4-mediated protein stabilization. These data suggest that CDK4 stabilizes Foxj1 through its kinase activity.

### CDK4 physically interacts with and directly phosphorylates Foxj1

To understand whether Foxj1 is a substrate of CDK4 kinase, we first examined the possible interaction between these two proteins. HeLa cells were transiently transfected with Flag-tagged zFoxj1a and Myc-tagged CDK4. Immunofluorescence staining experiments revealed a colocalization of overexpressed zFoxj1a and CDK4 in the nuclear aggregates (Fig 6A), suggesting a potential interaction between these two proteins. As zFox1a was observed to be localized exclusively to the nucleus, even when co-expressed with CDK4 (Fig 6A), we excluded the effects of CDK4 on the subcellular distribution of zFox1a. We next examined the association between zFoxj1 and CDK4 in wild-type embryos and HEK293T cells. Co-immunoprecipitation experiments demonstrated that overexpressed CDK4 interacted with endogenous and ectopically expressed zFoxj1a (Fig 6B and 6C). Interestingly, the kinase deficient form of CDK4 failed to associate with zFoxj1a (Fig 6C). In order to test whether CDK4 interacts directly with Foxj1 protein, we carried out an *in vitro* binding assay using purified proteins. As depicted in Fig 6D, Myc-CDK4 protein purified from bacterial cells was able to bind to GST-zFoxj1a but not GST proteins. Collectively, these results demonstrate that CDK4 interacts directly with zFoxj1a protein.

**Fig 6.**
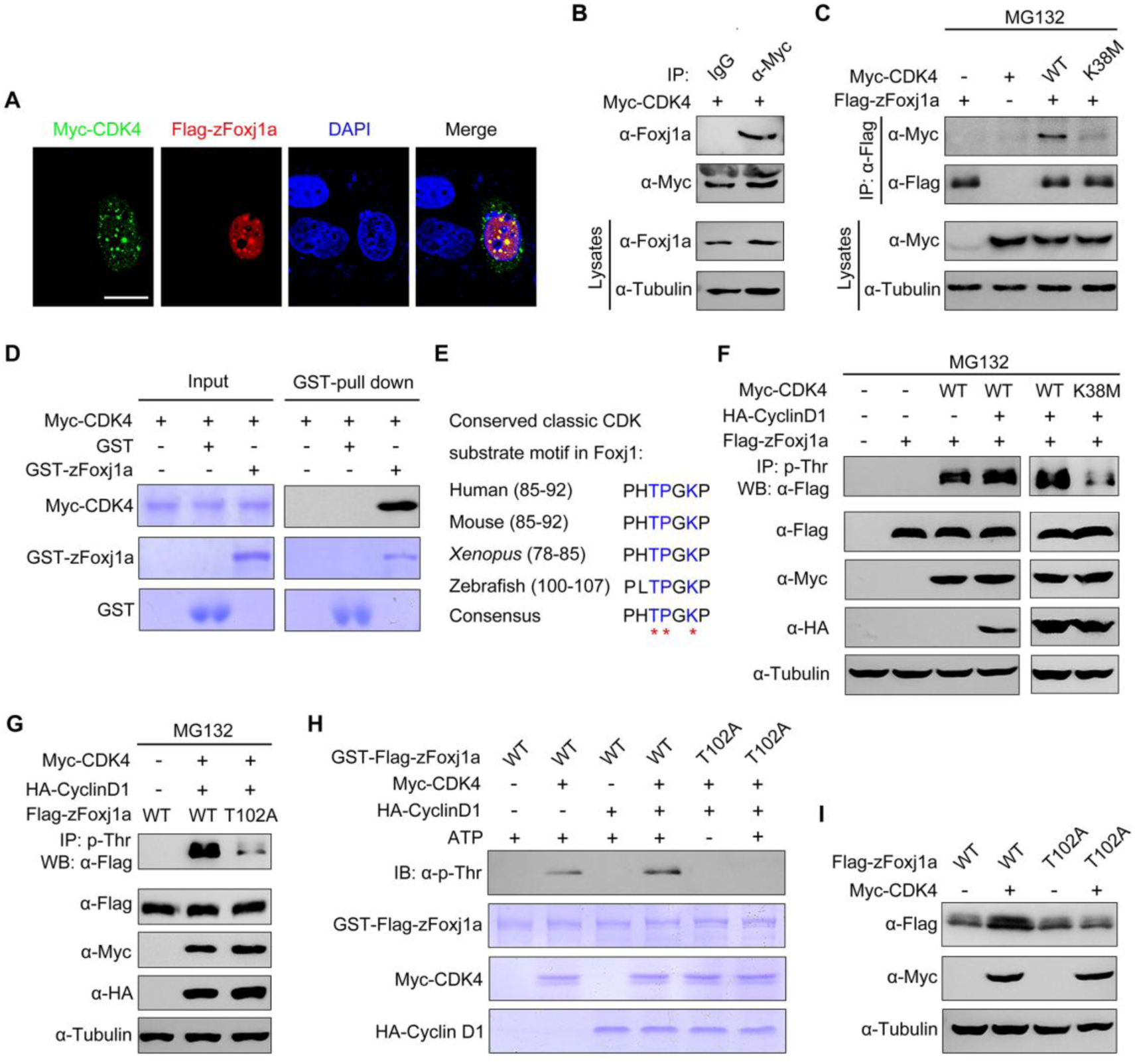
CDK4 phosphorylates zFoxj1a at T102 to suppress its degradation. (A) zFoxj1a and CDK4 show evident colocalization in Hela cells. Hela cells were transfected with Flag-zFoxj1a and Myc-CDK4 and immunostained with anti-Flag (red) and anti-Myc (green) antibodies. Nuclei were stained with DAPI (blue). Scale bar, 20 μm. (B) Overexpressed CDK4 interacts with endogenous zFoxj1a. Wild-type embryos were injected with 200 pg Myc-CDK4 mRNA at midgastrulation and then harvested at the bud stage for immunoprecipitation with anti-Myc antibody or normal mouse IgG. (C) CDK4 but not its kinase dead mutant interacts with zFoxj1a. HEK293T cells were transfected with plasmids as indicated, followed by treatment with MG132 for 5 hours prior to harvest for immunoprecipitation. (D) Direct binding of CDK4 to zFoxj1a *in vitro*. GST, GST- zFoxj1a and Myc-CDK4 were expressed in bacterial cells and purified. Myc-CDK4 proteins were incubated with GST or GST-zFoxj1a. The presence of Myc-CDK4 in the protein complex pull-downed by Glutathione agarose was assessed using an anti-Myc antibody. Input proteins were examined by Coomassie blue staining. (E) Conserved CDK substrate motifs in Foxj1 proteins from different species. Red stars indicate critical residues in the CDK substrate motifs. (F-G) CDK4 phosphorylates zFoxj1a at T102. HEK293T cells were transfected with the indicated plasmids. CDK substrates were immunoprecipitated using a phospho-threonine-proline antibody and blotted with anti-Flag antibody to detect phosphorylated zFoxj1a or zFoxj1a-T102A. zFoxj1a-T102A is an unphosphorylated form of zFoxj1a. Note that wild-type zFoxj1a could be phosphorylated by CDK4 but not by the CDK4-K38M mutant (F). CDK4-mediated phosphorylation was nearly abolished in zFoxj1a-T102A (G). (H) *In vitro* kinase assays revealed that CDK4 phosphorylates wild-type zFoxj1a but not the zFoxj1a-T102A mutant. zFoxj1a and zFoxj1a-T102A proteins were purified from bacterial cells and incubated with recombinant Cyclin D1 and CDK4 proteins in the presence or absence of ATP. Phosphorylation of zFoxj1a and zFoxj1a-T102A was detected by western blot using a phospho-threonine-proline antibody and input proteins were examined by Coomassie blue staining. (I) Ectopical CDK4 expression is unable to stabilize zFoxj1a-T102 mutant. Lysates from HEK293T cells transfected with the indicated plasmids were subjected to immunoblotting.

It is well established that the CDK families of serine/threonine protein kinases phosphorylate substrates containing the consensus amino acid sequence (S/T)PXR/K [49]. Because Cyclin D1-CDK4 complexes contribute to the stabilization of both mouse and zebrafish Foxj1 (Fig 5E and 5F), we hypothesized that CDK4 is involved in the phosphorylation of Foxj1 within conserved classic substrate motifs. Interestingly, we found that zFoxj1a contains a potential CDK phosphorylation motif “TPGK” at the N-terminal region, which is highly conserved in vertebrates (Fig 6E). To investigate whether CDK4 phosphorylates Foxj1 protein, a phospho-threonine-proline antibody was used to enrich CDK substrates from whole cell lysates, and the presence of phosphorylated Foxj1 was examined by western blot. With these experiments, we found that CDK4 or Cyclin D1-CDK4 complexes could effectively phosphorylate zFoxj1a and mFoxj1 (Fig 6F and S5A). As expected, CDK4-K38M almost lost the ability to induce zFoxj1a phosphorylation (Fig 6F). These results clearly indicate that Foxj1 can be phosphorylated by CDK4.

We then aimed to determine whether the threonine 102 residue (T102) within the putative conserved substrate motif of zFoxj1a is a major CDK phosphorylation site. Excitingly, we observed that Cyclin D1-CDK4 complexes significantly promoted wild-type zFoxj1a phosphorylation, which was nearly abolished in T102 mutant, an unphosphorylated form of zFoxj1a (Fig 6G). Similarly, *in vitro* phosphorylation assays showed that purified CDK4 or Cyclin D1-CDK4 complexes resulted in distinct phosphorylation events when incubated with recombinant wild-type zFoxj1a protein, but not T102 mutant (Fig 6H). In addition, we observed an increased expression of wild-type zFoxj1a while not its T102 mutant in HEK293T cells upon co-expression with CDK4 (Fig 6I). Similarly, CDK4 was able to control mFoxj1 stabilization through phosphorylation of the T87 residue, located in the conserved substrate motif (S5B-S5C Fig). Taken together, we showed that CDK4 directly phosphorylates Foxj1 to suppress its degradation.

### zFoxj1a undergoes Ubiquitin-independent proteasomal degradation via a direct interaction with Psmd4b

Previous studies have suggested a primary role of the ubiquitin-proteasome system in the elimination of abnormal proteins and selective destruction of regulatory proteins [50,51]. To explore whether ubiquitination is required for Foxj1 degradation, we examined the effect of Ub K48R/G76A overexpression on zFoxj1a degradation. If zFoxj1a degradation was determined to be ubiquitylation-dependent, we would expect zFoxj1a to be stabilized upon overexpression of Ub K48R/G76A, which serves as a dominant negative inhibitor of poly-Ub chain formation [52,53]. Indeed, the presence of Ub K48R/G76A but not wild-type Ub efficiently inhibited the turnover of β-catenin (S6A Fig), which would be phosphorylated by glycogen synthase kinase-3β and destined for ubiquitin-mediated degradation [54]. Unexpectedly, overexpression of Ub K48R/G76A was unable to promote the stabilization of Flag-tagged zFoxj1a (S6A Fig). Because the Flag epitope contains two lysine residues, we generated a lysine-free version of zFoxj1a, termed HA-zFoxj1a-K20R by replacing the Flag tag with the HA epitope and mutating all 20 lysine residues within zFoxj1a protein to arginine residues. We found that both the wild-type and lysine-less zFoxj1a were significantly stabilized with CDK4 overexpression (S6B Fig). Therefore, zFoxj1a is able to be degraded independently of ubiquitylation.

Increasing evidence suggests that a list of proteins which directly interact with proteasomal subunits are thought to be degraded through an ubiquitin-independent degradation mechanism [55]. It has been demonstrated that the 19S regulatory subunit Rpn10 plays a critical role in the recognition of ubiquitin-independent substrates [56–58]. Therefore, we examined whether zFoxj1a binds to zebrafish Psmd4b, the ortholog of mammalian Rpn10. Indeed, overexpressed or endogenous zFoxj1a was found to interact with Psmd4b (Fig 7A and 7B). Consistent with these observations, an *in vitro* binding assay revealed a direct binding between purified zFoxj1a and Psmd4b (Fig 7C).

**Fig 7.**
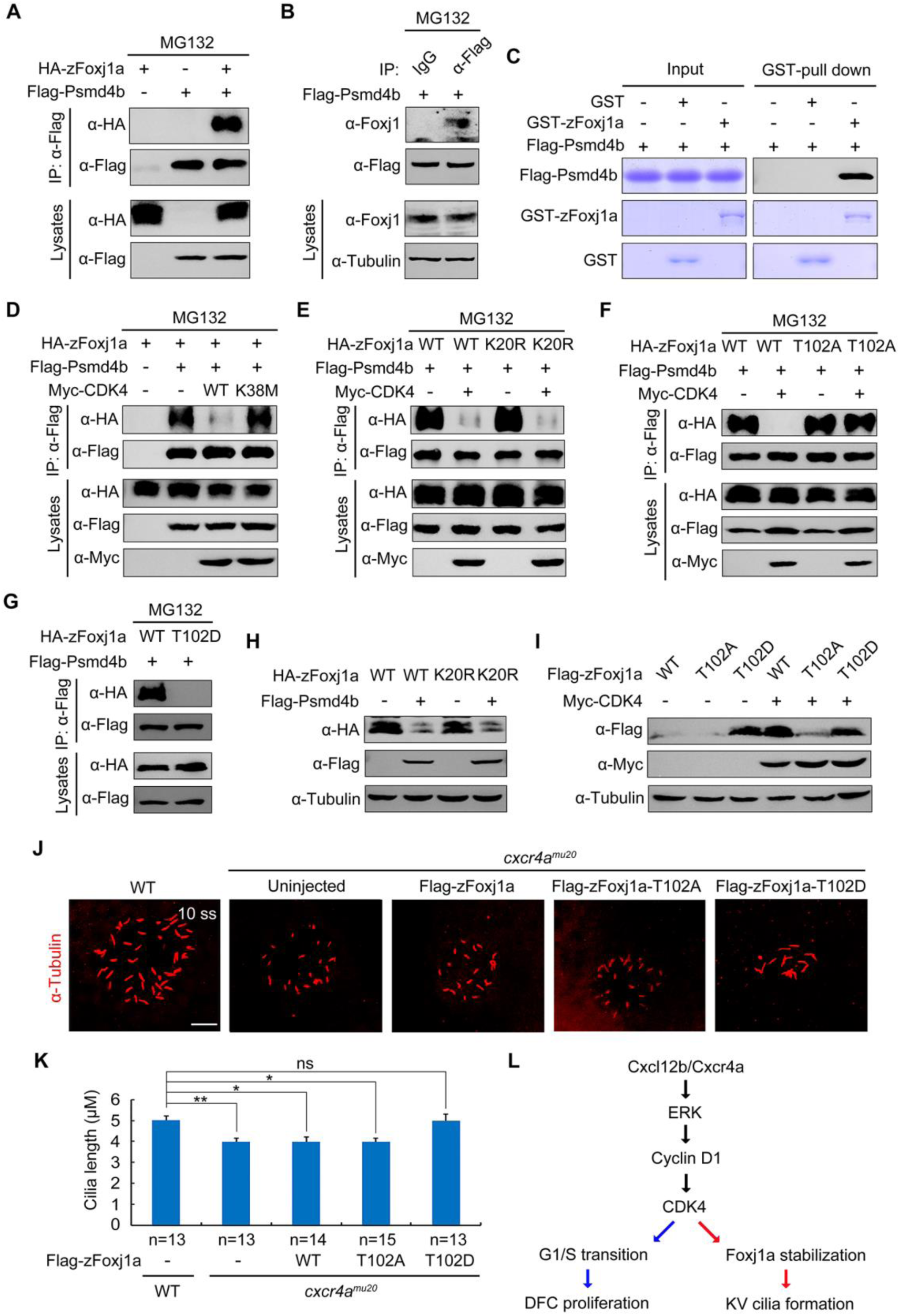
zFoxj1a undergoes Ubiquitin-independent proteasomal degradation via a direct interaction with Psmd4b. (A-B) Flag-Psmd4b interacts with overexpressed or endogenous zFoxj1a. HEK293T cells transfected with the indicated constructs (A) and bud-stage wild-type embryos with DFC-specific expression of Flag-Psmd4b (B) were subjected to immunoprecipitation with the indicated antibodies. (C) *In vitro* GST Pull-Down assays reveal a direct interaction between zFoxj1a and Psmd4b. Purified GST or GST-zFoxj1a proteins were incubated with recombinant Flag-Psmd4b. The presence of Flag-Psmd4b in the protein complex which was pull-downed by Glutathione agarose was assessed by western blot and input proteins were examined by Coomassie blue staining. (D-E) CDK4 kinase activity is required for its inhibitory effect on the association between Psmd4b and zFoxj1a. HEK293T cells transfected with the indicated plasmids were treated with MG132 for 5 hours prior to harvest for immunoprecipitation. Note that ectopically expressed CDK4 efficiently disrupted the association of Psmd4b with zFoxj1a (D) or its lysine-free mutant zFoxj1a-K20R (E), while overexpression of the CDK4 kinase deficient mutant CDK4-K38M had no effect on their interaction (D). (F-G) Psmd4b has a much lower affinity for zFoxj1a-T102D. zFoxj1a-T102A is an unphosphorylated form and zFoxj1a-T102D is a phospho-mimicking mutant of zFoxj1a. Note that Psmd4b was particularly associated with zFoxj1a and zFoxj1a-T102A (F), but was unable to bind zFoxj1a-T102D (G). The association between Psmd4b and zFoxj1a-T102A was unaffected by CDK4 overexpression (F). (H) Psmd4b overexpression induces a dramatic reduction in zFoxj1a expression. Note that a similar reduction in the expression level of the lysine-free mutant zFoxj1a-K20R was observed upon Psmd4b overexpression. (I) Comparison of protein stability in wild-type zFoxj1a and its mutants. HEK293T cells were transfected with the indicated constructs and harvested for western blot analysis. In comparison to wild-type zFoxj1a and its T102A mutant, the zFoxj1a-T102D mutant exhibited greater stability, but could not be further stabilized by CDK4 overexpression. (J-K) DFC-specific overexpression of zFoxj1a-T102D in *cxcr4a^um20^* mutants restores KV cilia length. Confocal images of 10-somite-stage *cxcr4a*-deficient embryos were injected with 200 pg of wild-type *zfoxj1a* or *zfoxj1a-T102A* or *zfoxj1a-T102D* mRNA at the 256-cell stage. The resulting embryos were harvested at the 10-somite stage for immunostaining using an antibody against acetylated tubulin (J). Scale bar, 20 μm. Statistical data for cilia length were shown in panel K. Student’s *t-*test, \**P*<0.05, \*\**P*<0.01. ns, no significant difference. (L) A schematic diagram showing the regulatory mechanism of Cxcl12/Cxcr4a signaling in DFC proliferation and cilia formation.

To further unveil how CDK4 and its kinase activity regulate Foxj1 stabilization, we examined whether CDK4 functions to control the interaction between Psmd4b and zFoxj1a. As expected, ectopic expression of wild-type CDK4, but not its kinase deficient mutant, dramatically suppressed the association of Psmd4b with zFoxj1a or zFoxj1a-K20R (Fig 7D and 7E). In contrast, the binding ability of the unphosphorylated form of zFoxj1a for Psmd4b was retained with CDK4 co-expression (Fig 7F). Interestingly, the phospho-mimicking mutant of zFoxj1a (zFoxj1a-T102D) completely lost its ability to bind Psmd4b (Fig 7G), suggesting that CDK4-mediated phosphorylation of zFoxj1a at T102 eliminates its affinity for Psmd4b. In addition, Psmd4b overexpression reduced the expression levels of zFoxj1a and its lysine-free mutant (Fig 7H). When HEK293T cells were transfected with same amount of plasmid DNA to express wild-type zFoxj1a and its T102A and T102D mutants, respectively, we detected the highest expression level of zFoxj1a-T102D (Fig 7I). However, T102D mutant could not be further stabilized by CDK4 overexpression (Fig 7I). Taken together, these results indicate that CDK4 phosphorylates and stabilizes zFoxj1a by disrupting its association with Psmd4b.

We next addressed the developmental relevance of CDK4-induced zFoxj1a stabilization. *Cxcr4a-deficient* DFCs exhibited defective *cycling D1* expression and impaired G1/S transition (Fig 3E-3H and 4F), implying a dysregulated activation of the CDK4/6 kinases. DFC-specific overexpression of zFoxj1a-T102D, but not wild-type or zFoxj1a-T102A, was found to restore the length of KV cilia in *cxcr4a^um20^* mutants (Fig 7J and 7K). As zFoxj1a is not involved in DFC proliferation [19,20], it was reasonable to find that the decrease in the number of cilia was not alleviated (S7 Fig). Overall, our results support a model in which Cxcl12b/Cxcr4a signaling activates ERK1/2, which then promotes Cyclin D1 expression. This in turn activates CDK4/6 kinase activity in DFCs. These activated G1 CDKs drive G1/S transition during DFC proliferation and promote zFoxj1a stability by phosphorylation to support KV ciliogenesis at later stages (Fig 7L).

## Discussion

In zebrafish embryos, DFCs undergo mitotic proliferation during epiboly and then exit the cell cycle, giving rise to epithelial cells that assemble cilia in the mature KV organ [16]. Cell cycle defects in DFCs are often accompanied by an alteration in KV cilia elongation, raising the issue of whether there exists a feasible link between the cell cycle and cilia formation [16]. In this study, our experiments resolve this issue by demonstrating that Cxcr4 signaling is required for DFC proliferation and KV ciliogenesis by promoting Cyclin D1 expression. Specifically, we found that Cyclin D1-CDK4/6 accelerates the G1/S transition in DFCs, while also facilitates cilia formation via stabilization of zFoxj1a. Ciliary dynamics appear to be precisely coordinated with cell cycle progression [59]. It has been suggested previously that cell quiescence is essential for the formation of mouse nodal cilia [60]. Indeed, we observed that proliferating DFCs enter into a quiescent state upon differentiation into ciliated epithelial KV cells. Interestingly, our data indicates that during epiboly stages, Cxcr4a signal-induced expression of Cyclin D1 functions to regulate DFC proliferation and zFoxj1a stability, which is important for the ciliogenesis of quiescent KV cells. Therefore, the rapid cell cycle progression of DFCs during epiboly stages is not only required for the generation of enough cells to construct KV, but also plays a critical role in reserving sufficient levels of zFoxj1a protein to support subsequent cilia formation. Because Wnt/β-catenin signaling has been reported to play a role in both DFC proliferation and KV cilia elongation [21,61], it is interesting to consider whether this signaling pathway contributes to zFoxj1a stabilization via regulation of cell cycle progression during the establishment of L-R asymmetry.

It has been reported previously that G1 cyclins function together with their associated CDKs to phosphorylate a variety of transcription factors, including Smad2/3 and pluripotency factors, to control embryonic stem (ES) cell differentiation [11,12]. A systematic screen for CDK4/6 substrates identified fox family transcription factor FOXM1 as a critical phosphorylation target [62]. CDK4/6 stabilize and activate FOXM1 by phosphorylation at multiple sites to protect cancer cells from senescence [62]. In contrast, CDK2 reduces DNA damage-induced cell death by phosphorylation of FOXO1 at Ser249, resulting in cytoplasmic localization of FOXO1 [63]. In this study, we show that CDK4 directly interacts with and phosphorylates zFoxj1a at a conserved “TPGK” motif within the N-terminal region. Phosphorylation at T102 was not found to alter the subcellular distribution of zFoxj1a, but was shown to promote its stabilization. Therefore, the functional interaction between CDK4 and zFoxj1a provides a mechanism by which cilia development is facilitated. Because CDK4 also stabilizes mFoxj1 through phosphorylation of T87 within the substrate motif, it is likely that the molecular linkage between cell-cycle progression and ciliogenesis is conserved among vertebrates.

The majority of proteosomal protein degradation relies on Ub conjugation. However, there are increasing numbers of examples of proteasomal degradation which occur without prior ubiquitination [64,65]. Our study reveals that overexpression of the dominant-negative Ub has no effect on zFoxj1a stabilization. However, wild-type and lysine-less zFoxj1a are found to be similarly stabilized by ectopic CDK4 expression. Therefore, zFoxj1a is targeted for proteasomal degradation in an Ub-independent manner. Intriguingly, E3 Ub ligases, including MGRN1 and Lnx2b, have been reported to play a role in L-R laterality specification in rodents and zebrafish [23,66], suggesting a role of the Ub-proteasome system in the modulation of protein turnover during L-R body patterning. However, due to the fact that L-R symmetry breaking occurs within a short time window during vertebrate embryonic development [4,67], the accelerated and economical regulation of protein degradation may be essential. Because Ub-independent degradation does not require the enzymatic cascade of Ub-conjugation, it would be more efficient to alter the concentration of zFoxj1a protein levels via Ub-independent proteasomal degradation during L-R asymmetric development. Interestingly, a recent study has demonstrated that Foxj1 is rapidly turned over by the Ub-proteasome system in mouse primary ependymal cells [68]. Therefore, Foxj1 is a protein with a short half-life which undergoes proteasomal degradation via Ub-dependent or -independent pathways dependent on the cellular context.

Several proteins have been reported to interact with the 19S regulatory subunit Rpn10 via their Ub-like (UBL) domains [56–58]. Interestingly, while lacking a UBL-domain, zFoxj1a interacts directly with Psmd4b, the zebrafish ortholog of mammalian Rpn10. Our study demonstrates that CDK4 phosphorylates and stabilizes zFoxj1a by disrupting its association with Psmd4b. Similarly, the Ub-independent proteasomal degradation of Yeast Pah1 has also shown to be governed by its phosphorylation state [69]. Therefore, this may represent a general mechanism by which protein kinase-mediated phosphorylation plays a critical role in the protection of their substrates from Ub-independent proteasomal degradation.

## Materials and methods

### Zebrafish strains

Wild-type embryos were obtained from natural matings of Tuebingen zebrafish. Embryos were raised in Holtfreter’s solution at 28.5 °C and staged by morphology. *cxcr4a* mutant embryos were generated by crossing homozygous male and female *cxcr4a^mu20^* adult mutants. *Tg(sox17:GFP)* transgenic embryos were used to indicate the DFCs and KV cells during L-R asymmetric development. *Tg(EF1α:mKO2-zCdt1(1/190))* transgenic embryos express the fluorescent fusion protein mKO2-zCdt1(1/190) in cells at the G1 phase during embryonic development. *Tg(flk:EGFP)* transgenic embryos express EGFP in blood vessels. Our zebrafish experiments were all approved and carried out in accordance with the Animal Care Committee at the Institute of Zoology, Chinese Academy of Sciences (Permission Number: IOZ-13048).

### RNA synthesis, morpholinos and microinjection

Capped mRNAs for *cxcr4a*, *caMEK1*, *cyclin D1*, *zfoxj1a, zfoxj1a-T102A* and *zfoxj1a-T102D* were *in vitro* synthesized from corresponding linearized plasmids using the mMessage mMachine kit (Ambion). Digoxigenin-UTP-labeled antisense RNA probes were in vitro transcribed using the MEGAscript Kit (Ambion) according to the manufacturer’s instructions. The standard control morpholino (5’-CCTCTTACCTCAGTTACAATTTATA-3’) and splicing MO targeting *cxcr4a* (5‘-AGACGATGTGTTCGTAATAAGCCAT-3’) were purchased from Gene Tools (Philomath, OR, USA) and used as previously described [30,70]. For DFC-specific knockdown or overexpression experiments, indicated MOs or mRNAs were injected into the yolk at the 256-cell stage as described previously [36].

### Whole-Mount in situ hybridization

Whole-mount *in situ* hybridization was performed using the NBT-BCIP substrate following standard procedures. For two-color fluorescence *in situ* hybridization, Anti-digoxigenin-POD (11633716001, Roche) and anti-fluorescein-POD (11426346910, Roche) were used as primary antibodies to detect digoxigenin-labeled sox17 probes and fluorescein-labeled *cyclin D1* probes, respectively. Fluorescence *in situ* hybridization was then carried out using the Perkin Elmer TAS fluorescein system (NEL701A001KT) according to the manufacturer’s instructions.

### Cell lines and transfection

HEK293T and Hela cell lines (American Tissue Culture Collection, ATCC, USA) were cultured in DMEM medium supplemented with 10% FBS in a 37°C humidified incubator in a 5% CO2 environment. Cell transfections were carried out using Lipofectamine 2000 (11668019, Invitrogen) following the manufacturer’s instructions.

### Immunostaining and confocal microscope

Embryos were fixed in 4% paraformaldehyde overnight. Fixed embryos were then rinsed with PBST for a total of 4 times every 5 minutes. Embryos were then blocked at room temperature for 1 hour in 10% heat-inactivated goat serum and then stained with the following affinity-purified primary antibodies overnight at 4°C: anti-β-Catenin antibody (1:500; ab6302, Abcam), anti-Cdh1 (1:200; GTX125890, GeneTex), anti-pERK1/2 (1:1000; 9101, Cell Signaling), anti-acetylated-Tubulin antibody (1:400; T6793, Sigma), anti-BrdU (1 1000; ab6326, Abcam), anti-α-PKC (1:200, sc-216, Santa Cruz), anti-GFP (1:1000; A-11122, Invitrogen), anti-GFP (1:1000; A-11120, Invitrogen), anti-Foxj1 (1:200; ab220028, abcam). Samples were then washed three times with PBST, followed by incubation with secondary antibodies, including DyLight 488-conjugated Goat anti-rabbit IgG (1:200; 711-545-152, Jackson), DyLight 594-conjugated Goat anti-mouse IgG (1:200; 715-585-150, Jackson), DyLight 488-conjugated AffiniPure goat anti-mouse IgG (1:200; 715-545-150, Jackson) and DyLight 594-conjugated AffiniPure goat anti-rabbit IgG (1:200; 711-585-152, Jackson) for 1 hour at room temperature. In some experiments, DAPI (1:10000, Sigma) was used to stain nuclei. The stained embryos were then embedded with 2% low melting agarose and imaged using a Nikon A1R+ confocal microscope with identical settings.

### Pharmacological treatment

To block CDK activity, embryos were treated with 0.5 μM PD0332991 (A8318, Palbociclib) or 0.2 μM CY202 (A1723, Palbociclib) from the shield stage to the bud stage. For CDK4/6 inhibition in cultured cells, HEK293T cells were treated with 0.5 μM PD0332991 for 5 hours prior to harvest. In order to examine which pathway is required for zFoxj1a degradation, HEK293T cells were transfected with plasmids expressing Flag-zFoxj1a and treated with 20 mM NH4Cl (A116363, Aladdin), 20 μM MG132 (M7449, Sigma) and 5 mM 3-MA (M9281, Sigma), respectively, for 5 hours prior to harvest.

### Fluorescent beads tracking

Fluorescent red beads of 1 uM diameter (1:500, 18660-5, Polysciences) were injected into the KV of embryos at the 6-somite stage. The resulting embryos were then embedded in 2% low melting agarose at the 10-somite stage for confocal imaging. Beads tracking videos and images were processed using Image Pro 6.0.

### Antibodies and immunoprecipitation assays

For immunoblotting experiments, we used the following affinity-purified antibodies: Anti-Flag (1:5000; F2555, Sigma), anti-Myc (1:3000; M047-3, MBL), anti-HA (1:3000; CW0092A, CW), anti-β-Tubulin (1:5000, CW0098M, CWBIO), and anti-Foxj1 (1:200; ab220028, Abcam).

For coimmunoprecipitation assays, embryos or HEK293T cells were harvested and lysed with TNE lysis buffer (10mM Tris-HCl, pH 7.5, 150 mM NaCl, 2 mM EDTA, and 0.5% Nonidet P-40) containing a protease inhibitor mixture. Lysates were incubated with anti-Flag-agarose beads (A2220, Sigma) or protein A-Sepharose beads (101041, Invitrogen) and anti-phospho-Threonine-Proline antibody (1:5000; 9391, Cell Signaling) at 4°C for 4 hours. Beads were washed four times with TNE buffer. Bound proteins were then separated by SDS-PAGE and visualized by western blots.

### In vitro GST Pull-Down

GST fusion proteins were expressed in Escherichia coli. strain BL21 and purified using Glutathione-Sepharose 4B beads (71024800-GE, GE Healthcare). GST-Myc-CDK4, GST-HA-cyclinD1, and GST-Flag-psmd4b were treated with Thrombin (1:1000; T4648, Sigma) to cleave their GST tags. For *in vitro* binding assays, GST-Foxj1a proteins were immobilized by Glutathione-Sepharose 4B beads and incubated with the indicated purified proteins at 4°C for 3 hours. Following washing, the bound proteins were separated with SDS-PAGE and analyzed by western blots.

### In vitro kinase assay

For i*n vitro* kinase assays, 1 μg GST-Foxj1a or GST-Foxj1a-T102A was incubated with 1 μg of the indicated purified proteins in 1×kinase buffer (25 mM Tris-Cl, pH7.5, 5 mM β-glycerophosphate, 0.1 mM Na3VO4, 10 mM MgCl2, 2 mM dithiothreitol) with or without 50 µM ATP (P0756S, New England Biolabs) at 30°C for 30 min. The mixture was then separated on 10% SDS-PAGE and visualized by western blots or Coomassie Blue staining.

### Statistical analysis

Cilia number and length were measured using ImageJ software. All results were expressed as the mean ± SD. Differences between control and treated groups were analyzed using the unpaired two-tailed Student’s *t*-test. Results were considered statistically significant at p <0.05.

## Acknowledgements

We are grateful to Dr. Jingwei Xiong (Peking University, China) for the *Tg(EF1α:mKO2-zCdt1(1/190))* fish line and Dr. Linfei Luo (Southwest University, China) for *cxcr4a^mu20^* and *cxcl12b^mu100^* fish lines.

## Supporting information

### Supplemental Figures

**S1 Fig.**
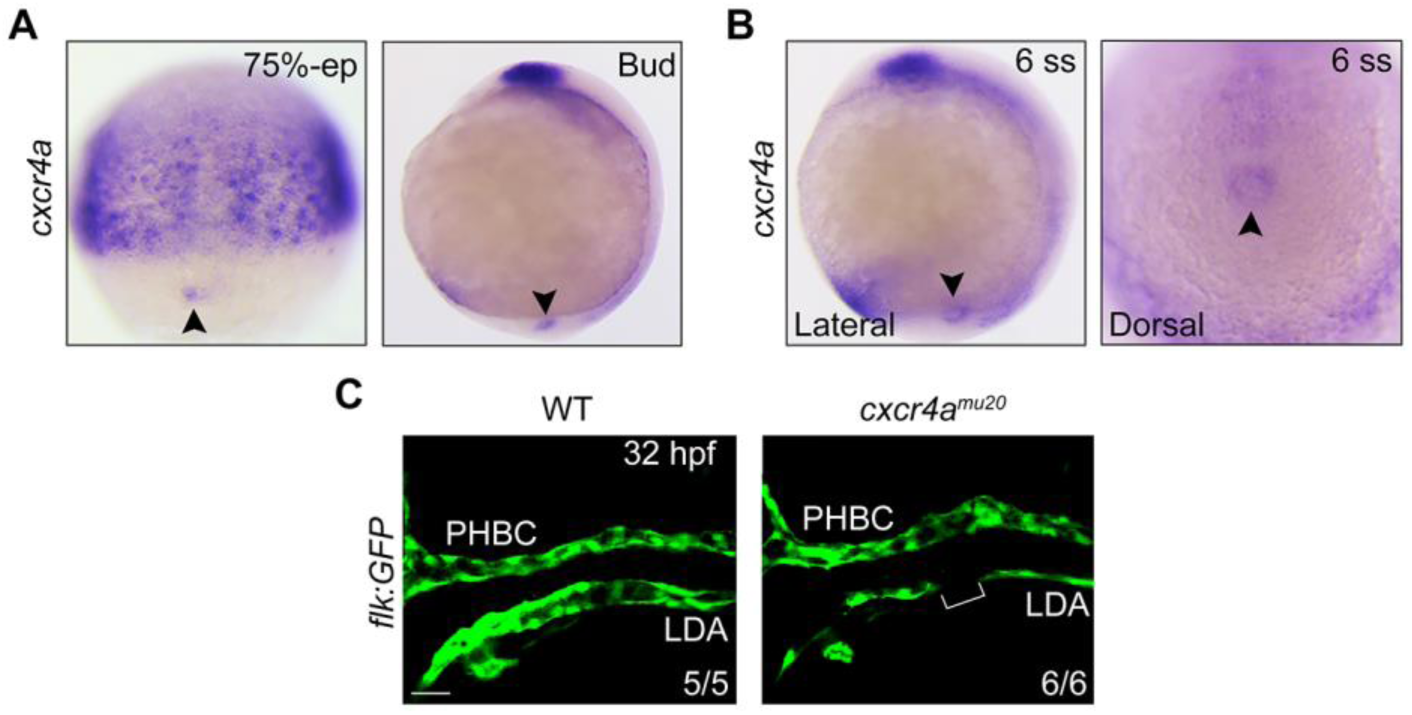
The expression of *cxcr4a* in DFCs and KV cells. (A) *cxcr4a* expression during gastrulation. *In situ* hybridization of *cxcr4a* in embryos at the 75%-epiboly stage (Dorsal view with animal pole to the top) and bud stage (Lateral views with animal pole to the top). Black arrowhead indicates the DFCs.75%-ep, 75%-epiboly. (B) *cxcr4a* expression at the 6-somite stage. Lateral view was shown with animal pole to the top in the left panel and dorsal view was shown in the right panel. Black arrowhead indicates the KV. (C) Confocal images depicting the formation of the lateral dorsal aorta in live *Tg(flk:GFP)* embryos. Scale bar, 50 μm. LDA, lateral dorsal aorta; PHBC, primordial hindbrain channel.

**S2 Fig.**
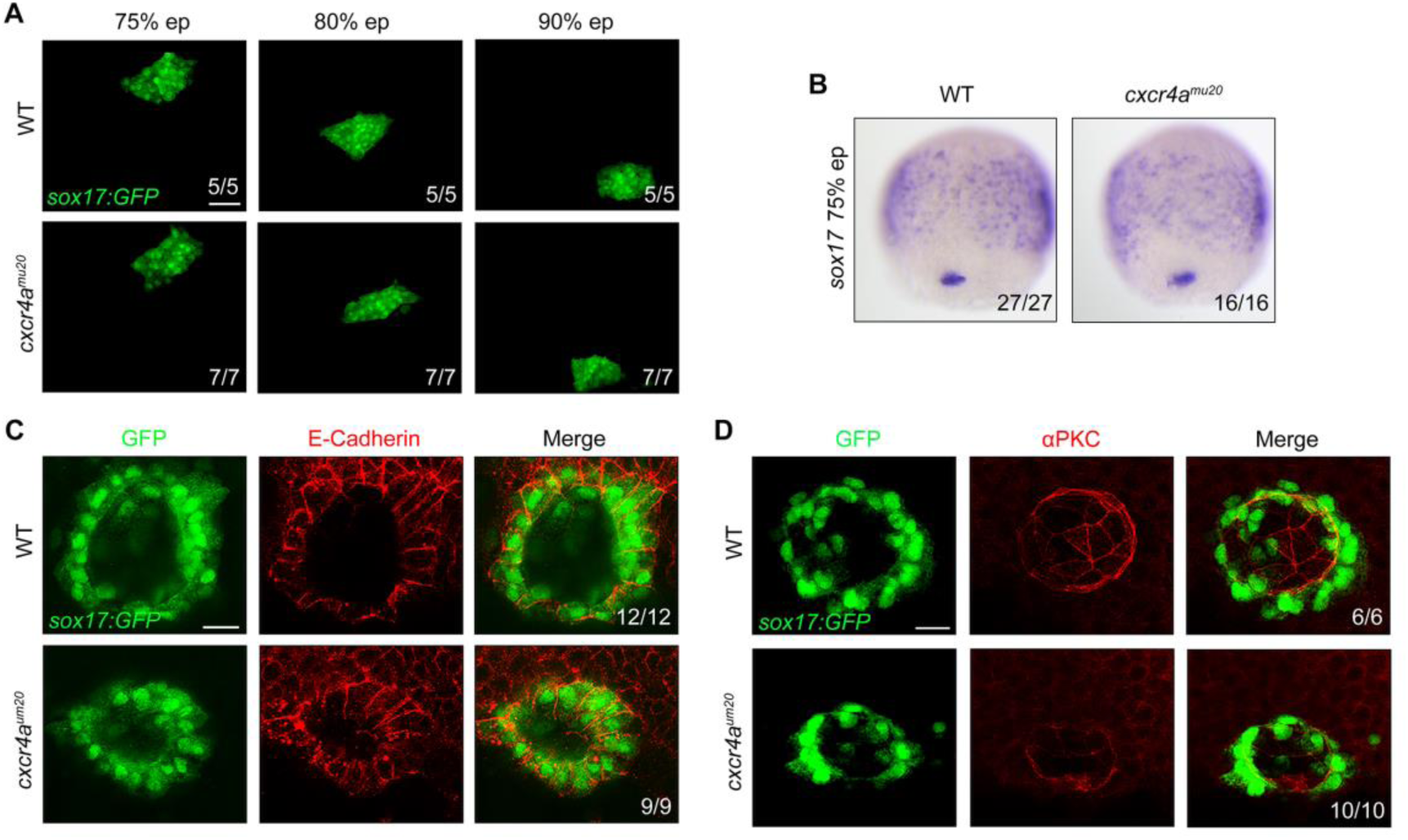
*cxcr4a* is unnecessary for the specification, clustering and collective migration of DFCs and dispensable for the polarized differentiation of KV cells. (A) Time-lapse confocal images showing DFCs migration in wild-type and *cxcr4a^mu20^* mutant embryos on a *Tg(sox17:GFP)* background from 75%- to 90%-eiboly stages. Scale bar, 50 μm. (B) Sox17 expression was examined by *in situ* hybridization in wild-type and *cxcr4a^mu20^* mutants at the 75%-epiboly stage. (C-D) Wild-type and *cxcr4a*-deficient *Tg(sox17:GFP)* embryos were harvested at the 10-somite stage for immunostaining. KV cells were labelled using an antibody against GFP (green). Expression of the basal-lateral marker E-cadherin (C) and the apical marker aPKC (D) were visualized using the indicated antibodies (red). Scale bar, 20 μm.

**S3 Fig.**
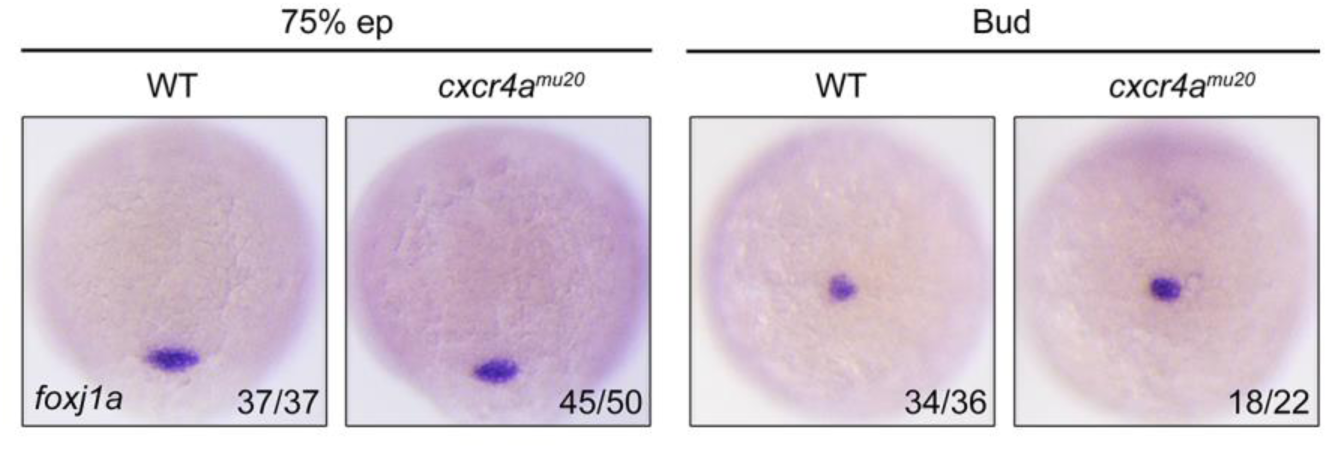
*cxcr4a^mu20^* mutants exhibit a normal expression of *cxcr4a* transcripts. *cxcr4a* expression was examined by *in situ* hybridization at the 75%-epiboly and bud stages in wild-type and *cxcr4a^mu20^* mutant embryos.

**S4 Fig.**
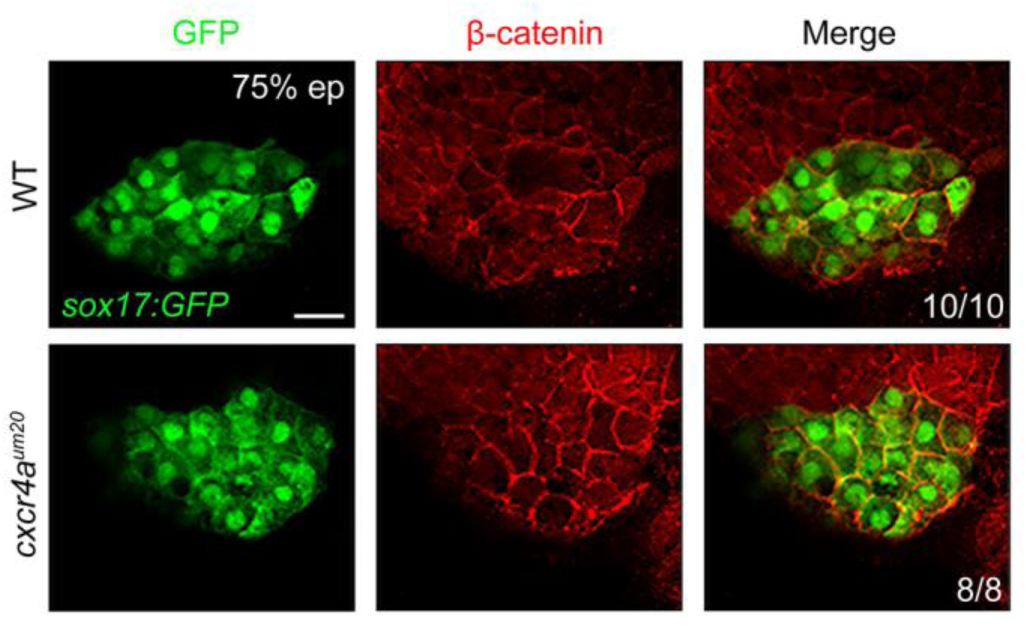
Inactivity of *cxcr4a* does not affect β-catenin nuclear accumulation in DFCs. Wild-type and *cxcr4a^mu20^* mutants were harvested at the 75%-epiboly stage for immunofluorescence assays using the indicated antibodies. Scale bar, 20 μm.

**S5 Fig.**
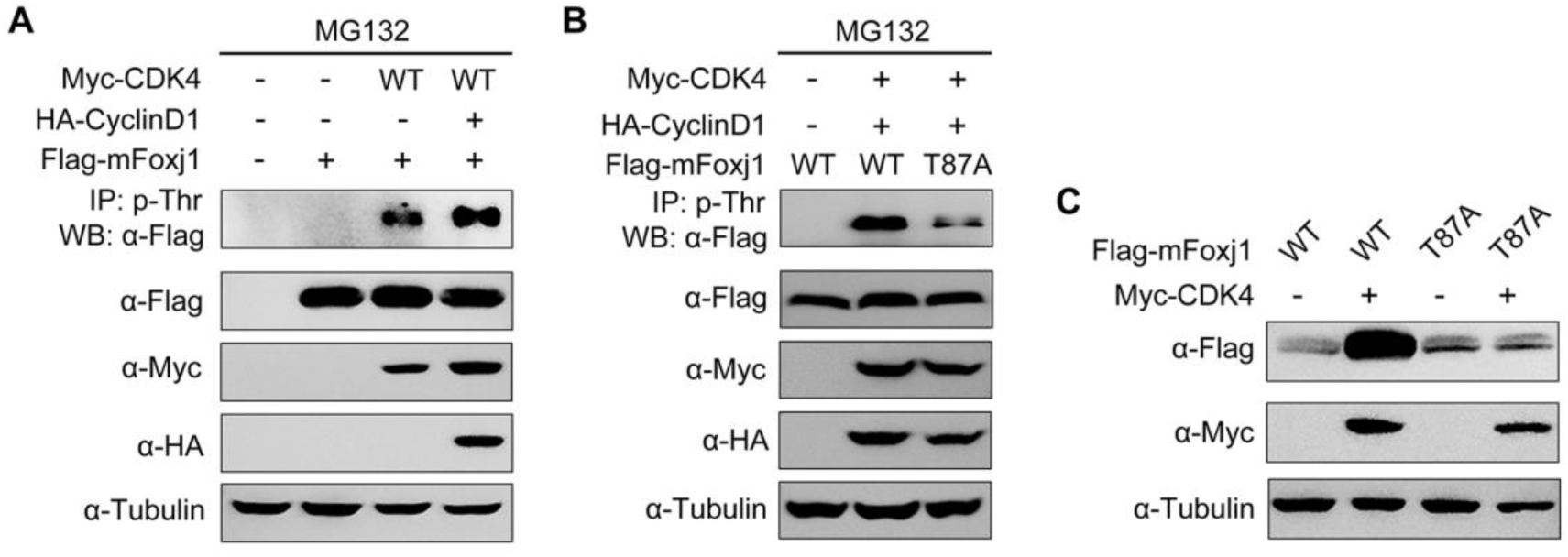
CDK4 phosphorylates and stabilizes mFoxj1. (A-B) HEK293T cells were transfected with the indicated plasmids and then harvested for immunoprecipitation with a phospho-threonine-proline antibody. Phosphorylation of mFoxj1 (A) and its T87A mutant (B) was detected by western blot. Note that the CDK4-mediated phosphorylation of mFoxj1 was clearly decreased in the T87A mutant. (C) Western blots of total lysates from HEK293T cells transfected with the indicated plasmids. Note that CDK4 overexpression could stabilizes wild-type mFoxj1 but not the T87A mutant.

**S6 Fig.**
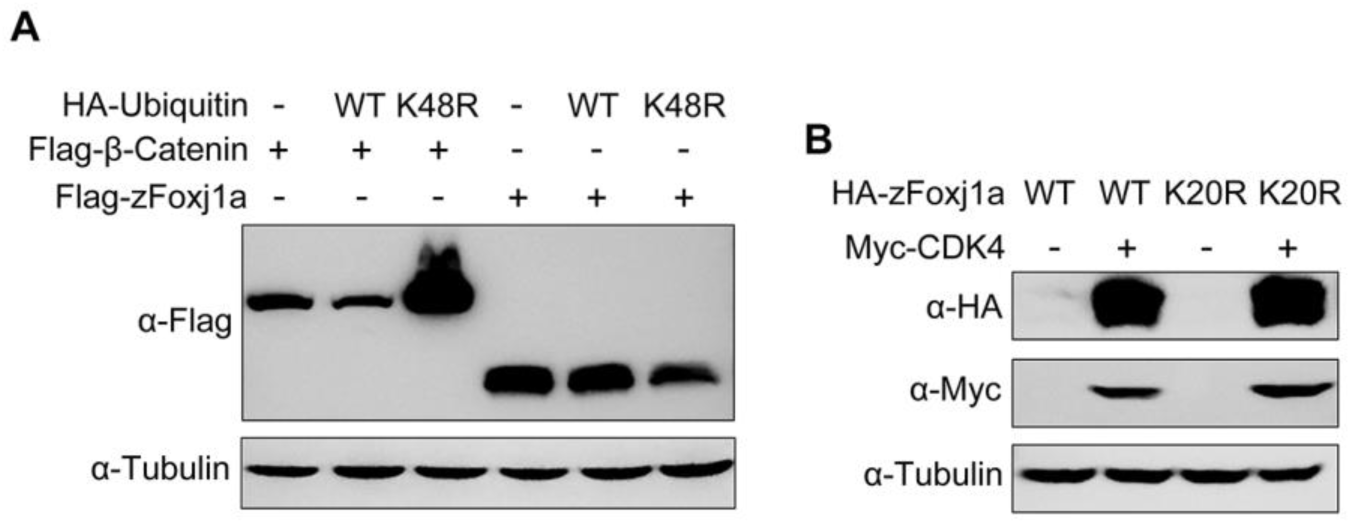
Proteasomal degradation of zFoxj1a is independent of ubiquitin modification. (A) Overexpression of Ub K48R/G76A was unable to stabilize zFoxj1a. Flag-tagged β-catenin and zFoxj1a were co-expressed with wild-type Ub or Ub K48R/G76A, a dominant negative inhibitor of chain formation and degradation. Cell extracts were immunoblotted with the indicated antibodies. (B) CDK4 overexpression similarly promoted the expression of wild-type zFoxj1a and its lysine-less mutant K20R.

**S7 Fig.**
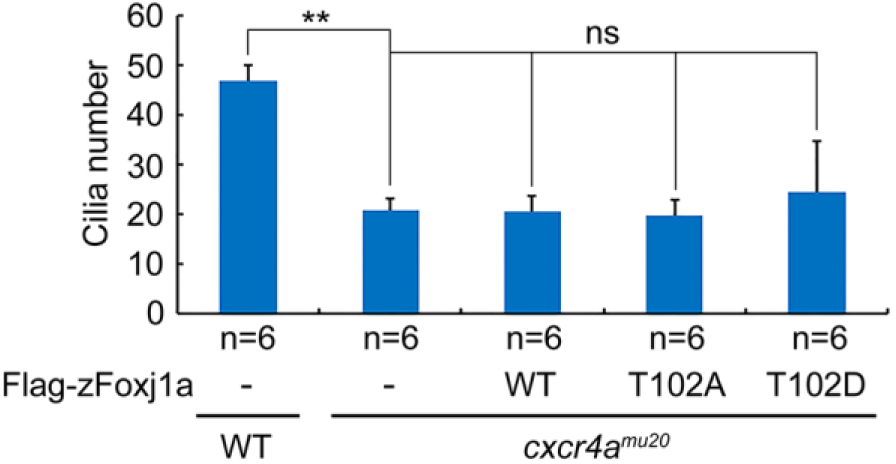
The decrease in the number of cilia in *cxcr4a^um20^* mutants was not alleviated by DFC-specific overexpression of wild-type zFoxj1a and its T102A and T102D mutants, respectively. *cxcr4a*-deficient embryos were injected with 200 pg of wild-type *zfoxj1a* or *zfoxj1a-T102A* or *zfoxj1a-T102D* mRNA at the 256-cell stage. The resulting embryos were harvested at the 10-somite stage for immunostaining using an antibody against acetylated tubulin. Cilia length was quantitatively analyzed using ImageJ software. Student’s *t-*test, \*\**P*<0.01. ns, no significant difference.

**S1 Video. Wild-type control embryos, normal KV flow.** Wild-type *Tg(sox17:GFP)* embryos were injected at the 6-somite stage with fluorescent beads and imaged using a Nikon A1R+ confocal microscope at the 10-somite stage. Dorsal view with anterior to the top.

**S2 Video. *cxcr4a^mu20^* mutant embryos, aberrant KV flow.** *cxcr4a^mu20^* mutant embryos on a *Tg(sox17:GFP)* background were injected at the 6-somite stage with fluorescent beads and imaged using a Nikon A1R+ confocal microscope at the 10-somite stage. Dorsal view with anterior to the top.

